# Acoustic Modulation of Ion‑Solvent Interactions: Microscopic Insights into Acoustoelectric Effect Modelling

**DOI:** 10.64898/2026.02.04.703740

**Authors:** Yuchen Tang, Wei Yi Oon, Wei-Ning Lee

## Abstract

The acoustoelectric (AE) effect, in which acoustic waves modulate the electrical properties of a conductive medium, holds significant potential for biomedical imaging. While classic models describe the phenomenon through conductivity modulation, a detailed understanding of its microscopic origins, particularly the role of ion behaviours, remains lacking. This study introduces a novel electrokinetic perspective by investigating how ultrasound modulates ion-solvent interactions, thereby bridging macroscopic AE signals with underlying ion dynamics. Through finite element simulations of a dilute NaCl solution, we demonstrate that acoustic pressure waves induce local variations in ion mobility and diffusion by altering ion hydration shells and solvent viscosity. These changes disrupt the balance among Coulombic, diffusive, and frictional forces on individual ions, leading to the local conductivity modulation. Furthermore, simulations reveal that acoustic perturbation of the electrode-electrolyte interface (EEI) significantly enhances AE signal generation, highlighting the EEI’s critical role in AE-related applications. By linking acoustic modulation to fundamental ion-solvent interactions, this work not only provides a foundation for more accurate, microscopically grounded models of the AE effect but also connects AE effect modelling to the active research of solvation dynamics in physical chemistry.

## Introduction

The acoustoelectric (AE) effect was first predicted by Debye [2] in 1933 that *periodic changing electric charge density will accompany the sound in an electrolyte*. Not until a study on a saline solution in 2000 was Debye’s prediction experimentally validated with ultrasound stimuli at 500kHz [3]. J. Jossinet and B. Lavandier et al. revisited the physics of the AE effect and modeled it in terms of the electrical impedance modulation from ultrasound [4]. Since then, the AE effect has been widely referred to as the local modulation of electrical conductivity in solution or biological tissues in the subsequent study with medical ultrasound. In the last two decades, the AE effect has gained increasing attention due to its potential biomedical applications, like Acoustoelectric Tomography (AET) [5], Ultrasound Current Source Density Imaging (UCSDI) [6-12], Ultrafast Acoustoelectric Imaging (UAI) [13, 14], and Acoustoelectric brain imaging (AEBI) [15-18] to measurement of bio-tissues’ electrical impedance or detect electrophysiological activities. Overall, the development medical imaging techniques based on the AE effect are progressing from the stage of “mathematical simulations and proof of concept on phantoms”[19] to the stage of “Proof of concept on animal organs”.

The acoustoelectric phenomenology modeled based on conductivity modulation by Jossinet et al. in the 90s [4, 20] laid the foundation of the aforementioned biomedical applications. Adapted from the literature, Figure 1 shows that the involved acoustoelectric effects in a solution can be categorized into three groups:

1. The molarity changes due to the adiabatic compression;
2. The changes in ion mobility due to the changes in solvent viscosity against pressure and temperature;
3. The changes in dissociation equilibrium of the partially dissociated species.

**Figure 1.**
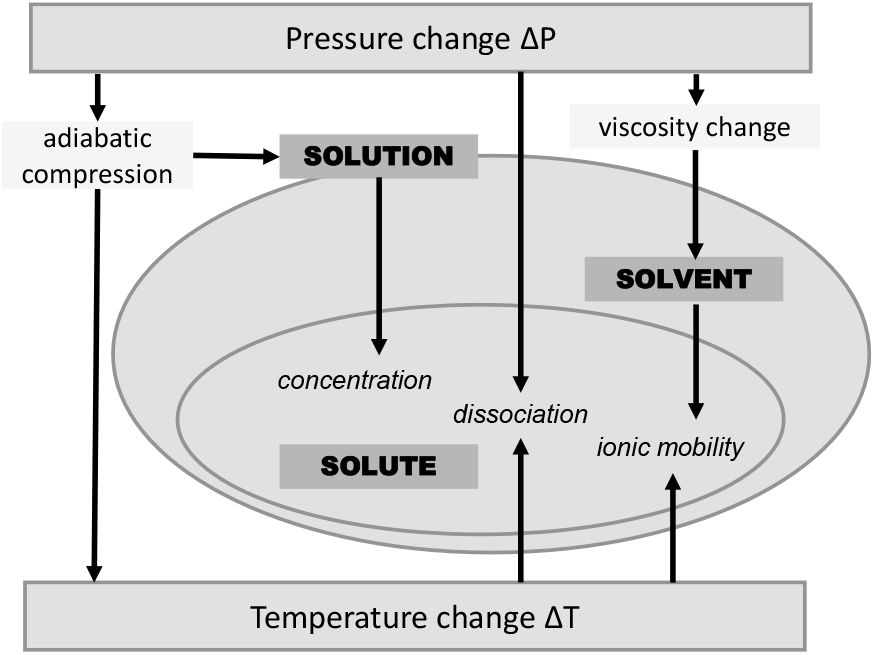
Graphic illustration of the mechanisms responsible for US modulation of the electrical conductivity in electrolytes. This figure is adapted from [4].

The sum of the three terms, including the bulk compressibility, the pressure coefficient of the solvent viscosity, and the change of the ion mobility against pressure, was contracted to an interaction coefficient, *K*(Unit: %/MPa) [4] as follows:

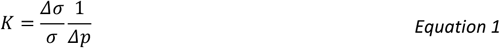

*K* was widely used in subsequent works to quantify the electrical conductivity/resistivity changes in response to ultrasound modulation [6, 21-23]. The AE interaction coefficient *K* was experimentally assessed in a NaCl solution, a CuSO_4_ solution, and a rabbit cardiac tissue by [9] based on the conductance changes at the US focal zone. The interaction coefficient *K* provides a solid interpretation of the AE effect from the macroscopic perspective. However, as it collectively quantifies the three modulated terms in the AE effect and could be only measured from a macro view, *K* cannot reveal how these terms are individually modulated by acoustic waves. This limitation is particularly evident in the case of ion mobility, which is derived from ion migration at the microscopic level, while conductivity is derived from the conduction of the bulk medium measured on the macroscopic view. Establishing a link between macro measurements and micro quantifications could offer a full picture and enhance our understanding of the inherent physics of AE effect.

Besides the theoretical gap, there are discrepancies in experimental findings due to the weak signal magnitudes (∼µV). The imaging techniques based on the AE effect generally have a low Signal-to-noise Ratio (SNR) and are easily perturbated by the parasite effects under certain experiment conditions. The identification of these parasitic effects varies depending on the various definitions of “AE signals” and “signals of interest” used in different studies, complicating the interpretation of results. For instance, back in 1950s, AE-like signals can be experimentally acquired when ultrasound presents even in a “pure liquid”, such as distilled water [24], methanol and ethanol [25], where the electrode-glass-solution interface may contribute to spurious responses. Despite decades of research, the origin of these AE signals remains unclear, as noted in the phenomenological study in 1990s [20]. The experimental measurement in 2000 [3] showed that the maximum signal was generated when the angle between the electric current direction and the ultrasound propagation direction was 80°, not theoretically-believed 90°. They pointed out that the measurement probably contained more than one signal component that had never been detected. Due to the scarce theory, these unanticipated signal components are hard to identify, whether as artifacts to be eliminated or as parasite effect that partially contribute to the “AE signals” of interest.

Further complicating the picture, classic AE phenomenology [2, 4] predicted the modulated potential should be at the insonification frequency, whereas recent studies showed that the generated signals exhibited a frequency that matched the pulse repetition frequency (PRF) of 100-1kHz from the 1 MHz insonification. Although this discrepancy lacks a theoretical explanation, the measured signals at PRF formed the basis of the Acoustoelectric Brain Imaging [17] and were used to effectively map the Steady State Visually Evoked Potential (SSVEP) of living rats [18]. Another inconsistency arises from the theoretical prediction from [4] concluded that the AE effect should be largely independent from on the ion species in terms of conductivity modulation, whereas the experimentally measured AE interaction constant *K* for NaCl and CuSO_4_ solution significantly differed [9]. The underlying physical mechanism is puzzling and remains to be explored.

This work aims to revisit the fundamental physical mechanisms and to understand the inherent processes and phenomena that lead to the generation of AE signals in various conditions through computational simulations. To achieve this, we first review theoretical foundation of AE phenomena, beginning with the relationship between electrolyte conductivity and ion mobility. Next, we will examine the acoustic modulation of electrolytes by incorporating thermodynamic potential and ion-solvent interactions, refining the theoretical framework for more accurate AE modeling.

## Methods

### From Macro to Micro: Conductivity to Ion Behaviour

The electrical conductivity κ(kappa, SI unit: S/m) is defined as *the conductance of a cube with an edge length measured at the right angle to the plane of the cube* [26]:

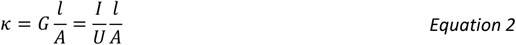

where *G* is the electrical conductance (SI unit: S), which quantifies the capability of an electrical conductor to carry an electric current via the Ohm’s law: 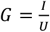 (*I*: current and *U*: voltage) *l, A* are the length and surface area of the conductor, respectively.

Ion mobility *u* (SI unit: m^2^.s^-1^.V^-1^) is defined as the absolute value of the ion velocity in an electrolyte per unit electrical potential difference

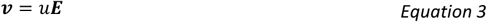

where *v* is the ion velocity (SI unit: m/s), ***E*** is the electrical field (SI unit: V/m).

To connect the measurable macro-scale property of conductivity with micro-scale ion characteristics, the bridging concept is the electrical current density ***J***(SI unit: A/m^2^). Macroscopically, ***J*** is written as the product of conductivity and the electrical field ***E***; while microscopically, it is modeled via ion velocity *v* as the charge flux of one ion species through the surface with unit area [27]:

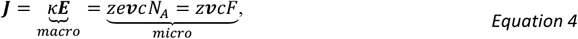

where the *z* is valence value; *c* is ion molarity, *e* is the elementary charge; *N*_*A*_ is the Avogadro constant (numbers of constituent particles per mole); *F* is the Faraday constant (*F* = *eN*_*A*_). Combining, Equation 3 and Equation 4, we obtain:

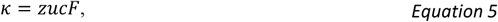

which links conductivity κ to ion mobility *u*. Considering multiple ion species in the electrolyte, the equation above can be rewritten as:

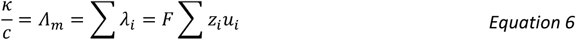

where Λ_*m*_ is the molar conductivity, *λ* is the ionic conductivity, and *i* is the number is the ion species. Molar conductivity Λ_*m*_ describes the ability of a solute to conduct an electric current by taking into account all the ions present [28], alternatively referring to as *molar conductance* and *equivalent conductance* in some literatures. Ionic conductivity *λ* characterizes ion migration in the solution for each species.

Direct experimental measurement of individual ionic conductivity *λ*_*i*_ is inherently limited, as the molar conductivity is not species-selective. Individual ionic conductivities *λ*_*i*_ can be calculated only if the ionic conductivity of one ion is known and the solution components are simple, containing only one cation and one anion. In complex mixtures, isolating each ion’s contribution is extremely difficult [27], because *λ*_*i*_ depends on numerous factors, such as concentration, ion mobility, solvent properties, temperature, and pressure, etc. In addition, *λ*_*i*_ varies with the solution composition due to different ion-ion interactions. Therefore, while direct experimental measurement remains difficult based on current instrumentation, a complete and accurate understanding of individual ionic conductivity can be achieved by complementing experiments with theoretical analysis of ion behaviors.

### Acoustic Modulation of Thermodynamic Energy

In physics, *phase* refers to a form of matter that is homogeneous in both chemical composition and physical state. To quantify the ability of a given phase component to change its state, *chemical potential μ* (the molal state function) is defined as the partial derivative of the thermodynamic potential (Gibbs free energy) with respect to the amount of substance *N*_*i*_ of a species *i*. The relationship between the chemical potential changes and the natural variables of a thermodynamic system are formulated as the Gibbs-Duhem equation:

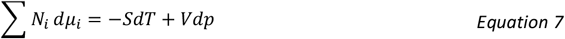

where *T, p, S* and *V* refers to temperature, pressure, entropy and volume, respectively.

For a phase in equilibrium, uniformity in the chemical potential is observed across all regions. Any disparity in chemical potential prompts the movement of system components from regions of higher chemical potential toward regions of lower chemical potential. When two phases with charged particles are brought into contact, the equilibrium between the two phases is established through the equalization of *electrochemical potential*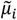:

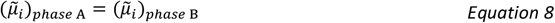

The electrochemical potential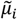 is defined as the sum of the chemical potential and the electrical work associated with a transfer of the charge *z*_*i*_*F* from infinity to the given place with electrical potential *ϕ*

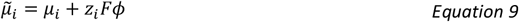

In other words, the electrical potential difference rises within the interphase to compensate for the chemical potential differences of the two phases. Thus, the equalization of the electrochemistry potential between the two phases in Equation 8 can be further written as

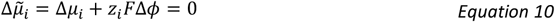

For example, when immerging a metal electrode into an electrolyte, the electrical potential rises in the vicinity of the electrode surface, forming an *electrical double layer (EDL)* to compensate for the chemical potential difference between the metal electrode and the electrolyte. The EDL encompasses the Stern layer (Helmholtz layer) and the Diffuse layer. The Stern layer is only several nanometers in thickness, and the particles there are influenced by the intermolecular force. The thickness of the EDL is dominated by the thickness of the diffuse layer, which is characterized by the Debye length *λ*_*D*_:

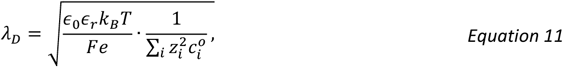

which is derived from the Poisson-Boltzmann equation, since the particles are mainly driven by Coulombic force and follow a Boltzmann distribution.

This principle of double-layer formation extends beyond static electrode interfaces. When an ultrasound wave propagates through the electrolyte, its alternating compressional and rarefactional phases act on the electrolyte along the way and form the cascaded liquid-liquid interphases. Within each phase, the ultrasound wave changes the thermodynamic equilibrium of the electrolyte via the local modulation of the pressure, temperature, and volume, resulting in variations of the local chemical potential (Equation 7). Consequently, the periodic ultrasound modulation of the local chemical potential generates a vibrational electrical potential in the interphase region. The series of separated layers can be regarded as a series of double layers [29] that compensate for the chemical potential differences between the alternating compressional and rarefactional phases of the ultrasound wave. The series of double layers can be regarded as a kind of “acoustic double layer”. When there is a pre-existed electrical field, the original electric current flow is affected by the formation of the “acoustic double layers”, thereby causing local acceleration or deceleration of the charges in different acoustic phases. This leads to a conductivity modulation of the medium in a macroscopic view.

To gain a comprehensive understanding of the local conductivity modulation from the formation of “acoustic double layers”, a detailed investigation of ion behavior within each acoustic phase is essential. The collective response of ions to this periodic modulation governs the net electric current flow. This principle can be illustrated with an analogy. consider a motorcade is delivering goods through a forest. The convoy’s overall efficiency depends not only on each vehicle’s capacity but also on the local condition of the forest, like terrain density and weather. As Figure 2 shows, an ultrasound wave modulates the electrolyte in a manner analogous to these changing environmental conditions. To predict the motorcade’s efficiency, one must understand how each vehicle reacts to hills, thickets, or storms. Similarly, to predict the electrolyte’s conductivity, we must understand how each ion’s migration changes as it travels through the alternating compressional and rarefactional phases of the ultrasound wave in response to the periodically varying chemical potentials. This transition from chemical potential to ion migration shifts our understanding of the AE effect from thermodynamics to electrochemistry, suggesting a focus on the influence of thermodynamic energy on individual ion behaviors.

**Figure 2.**
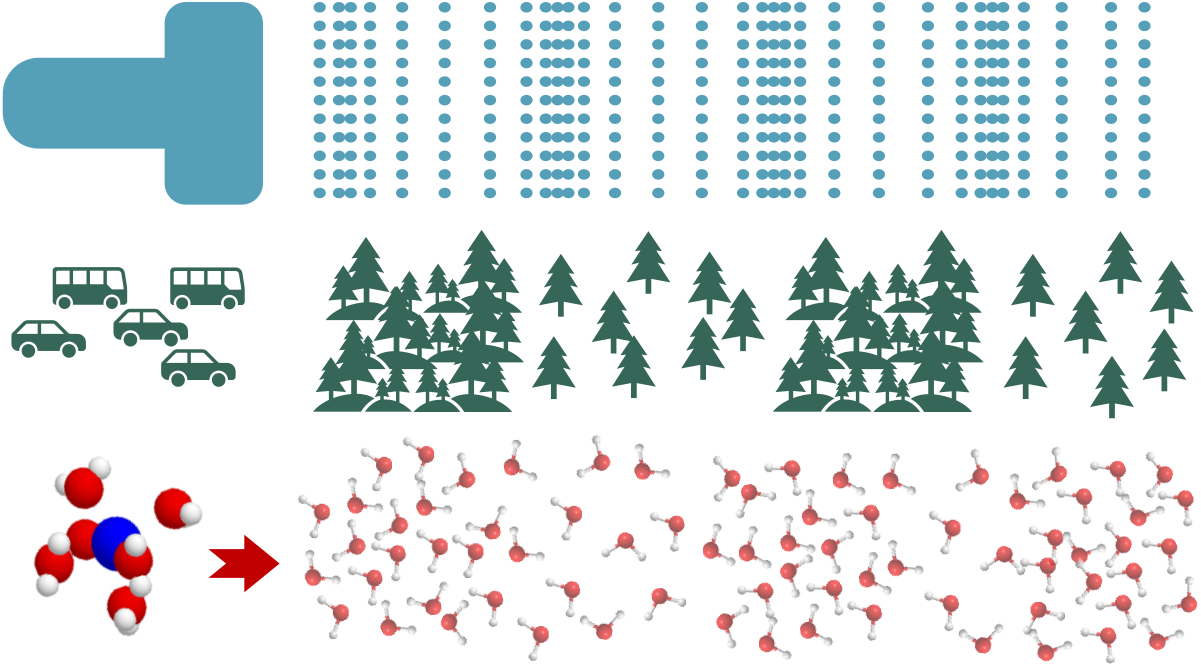
An illustration of how ions travel through the different phases under acoustic modulation. Cars and buses represent different ion species. The regions with different forest densities are analogous to the acoustic modulation phases.

### Acoustic Modulation of Ion-solvent Interaction

To model this individual ion behavior, a key physicochemical factor must be considered: ion hydration or solvation. The ion and its ‘backpack’, the tightly bound shell of solvent molecules, migrates as a cohesive unit. In the case of an aqueous solution, the water molecules orient themselves around the solute ions, forming the *Hydration Shell* (or *Solvation Shell* for other solvents).

An interchangeable concept is *Ion Atmosphere* which was proposed by Peter Debye and Erich Hückel in 1923 [30]. They pointed out that ions in solution are surrounded by a cloud of solvent molecules along with other counter ions, forming the ion atmosphere. Generally speaking, ion atmosphere encompasses a broader range of ions and solvent molecules attracted to the central ion, while the solvation shell specifically refers to solvent molecules.

The thickness and composition of the ion atmosphere depend on various factors, including the specific ion and solvent species, concentration, and natural variables like pressure and temperature. Among these, pressure, in particular, plays a particularly significant role by directly compressing or expanding the solvent matrix. As pressure increases, the solvent molecules packed more densely around the ion, strengthening electrostatic ion-solvent and thereby altering the stability, size, and organization of the hydration shell. Figure 3 briefly compares the chlorine hydration shell under normal (left panel: 0.1 MPa) and high (right panel: 4 GPa) pressure [31]. The water molecule numbers (hydration number or coordinate number), the radius, and the structure of the hydration shell alter due to the pressure changes.

**Figure 3.**
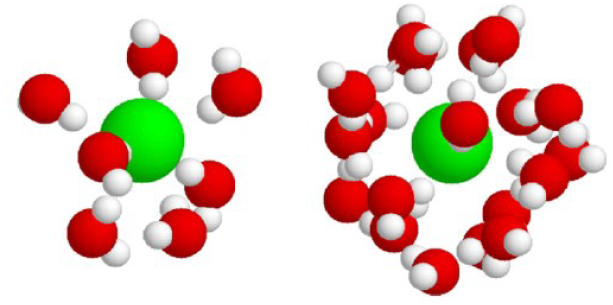
The chlorine hydration shell under normal and high pressures. The figure is adapted from [31].

The ion atmosphere critically influences both the physical properties of a solution and the dynamics of ion transport. For instance, it reduces the solution’s compressibility compared to pure water, because the water dipoles within the ion atmosphere are already partially oriented and compressed by the electric field of the central ion, making them less responsive to further compression compared to the ‘free’ water molecules in pure water. Moreover, ion atmosphere is integral to ion migration. Ions do not move solely but travel together with their surrounding solvation shell and diffuse ion cloud. In terms of fluid dynamics, the effective size of this migrating unit is described by the *Stokes Radius* (or *Stokes–Einstein Radius*). The radius determines the *Frictional Force* of the moving ion cloud based on Stokes law:

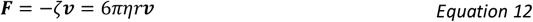

where ζ is the frictional coefficient; η is the solvent dynamic viscosity (SI unit: kg m^−1^ s^−1^); *v* is the speed of the spherical particle

Building directly on the hydrodynamic role of the Stokes radius, its quantitative impact on ion transport is captured by two fundamental relationships: the Stokes-Einstein law of diffusion (Equation 13) links the *Diffusion Coefficient D* to ion solvation shell radius *r*, while the Einstein–Smoluchowski (Equation 14) connects the ion mobility *u* directly to the frictional coefficient ζ which is a function of *r*.

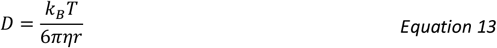

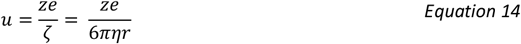

Collectively, these equations show that the effective solvation shell radius *r* directly governs both the ion’s mobility *u* and its diffusion coefficient *D*. The Stokes-Einstein equation (Equation 13) serves as a starting point for many research works in bridging the macroscopic system and molecular levels [32], while the Einstein–Smoluchowski (Equation 14) explicitly reveals how mobility is inversely proportional to frictional drag [27].

A second critical parameter in these equations is the solvent viscosity *η*, which is also pressure dependent. When regarded as an entity, dynamic viscosity *η* denotes the measurement of a fluid’s resistance to shear stress; when describing a solvent, the dynamic viscosity quantifies the internal frictions of the substrate. Liquids are assumed to be incompressible with pressure-independent viscosity in most research fields. However, in precision contexts of acoustic-electrical signals at the microvolt scale, even pressure-induced changes in *η* could become significant. These subtle variations can measurably affect ion mobility and diffusion, thereby contributing to the generation of the AE signals.

Building on the framework above, we hypothesize that acoustic modulation of the ion-solvent interaction is one underlying mechanism of the AE effect. This modulation operates on two fronts:

1. Modulation of the ion hydration shell radius on the ion side;

2. Modulation of the solvent viscosity on the solvent side.

As illustrated in Figure 4, such modulation disrupts the dynamic equilibrium among the diffusive, Coulombic, and frictional force acting on each ion cloud. The resulting imbalance induces local variations in the ion diffusion coefficient and ion mobility, which are the fundamental migration properties for charge transport. These microscopic perturbations propagate through the electrolyte as formulized by the Nernst-Planck equation, manifesting as measurable alterations in charge flow that generate the observed AE signals.

**Figure 4.**
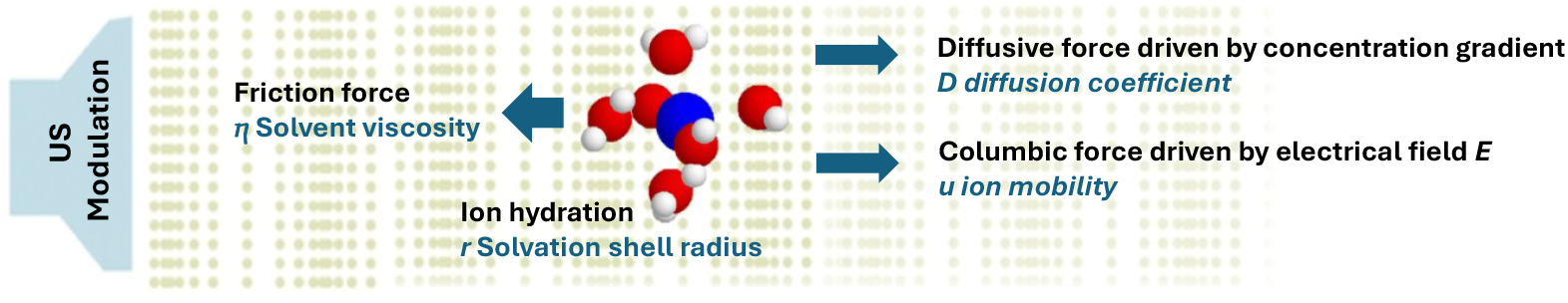
An illustration of the study hypothesis. For each ion cloud, three main forces are considered to be balanced: the diffusive force driven by the concentration gradient, the Coulombic force driven by the electrical field, and the frictional force.

Nevertheless, quantitatively modeling this pressure-modulated ion–solvent interaction remains challenging. There is no well-established formula that models an ion hydration structure under different pressures. It is still an active topic to investigate the pressure dependency of the H-bond [33] and O-bond [34] in the field of molecular dynamics, and the pressure dependency of the *Radial Distribution Function* of hydration for different ion species in the field of statistical mechanics [31, 35]. Similarly, the viscosity theory of liquids is still inadequately developed due to the complicated intermolecular forces [32, 36]. Current approaches often rely on empirical equations fitted to experimental measurements of the solution viscosity under different pressures [37].

Despite the incomplete theory for pressure-dependency of the ion-solvent interaction, a pilot simulation work can provide insight into the ion behavior underlying the AE effect, extending our interpretation from acoustic modulation on conductivity to a microscopic level. This new perspective on AE effect generation may potentially elucidate unexpected experimental and bridge theoretical advancements in AE-based functional ultrasound imaging to the ongoing research topics in chemical physics, thereby providing a promising direction for the development of AE theory.

### Finite Element Modelling

To investigate this hypothesis regarding the acoustic modulation of ion-solvent interactions, we conducted a finite-element simulation of a representative AE field. Figure 5 shows the simulation domain of a 2D conductive medium (29.6×14.8 mm: 50×25 ultrasound wavelength) containing a dilute NaCl solution (0.001 mol/m^3^) with COMSOL Multiphysics®. Two platinum electrodes (diameter: 0.6 mm, standard electrode potential: 1 V) were immersed in the medium, providing a voltage source of +500 mV or -500mV. A plane-wave ultrasound (US) pressure field (2.5 MHz, 2 cycles, 2.5 MPa) was simulated using the k-Wave® toolbox and imported into the FE model. Two configurations (Figure 5) were studied: ultrasound propagation parallel to the lead vector (S+ to SGND) in configuration A (left panel), and perpendicular in configuration B (right panel). The resulting AE signals were exported to MATLAB® 2022b for further signal analysis.

**Figure 5.**
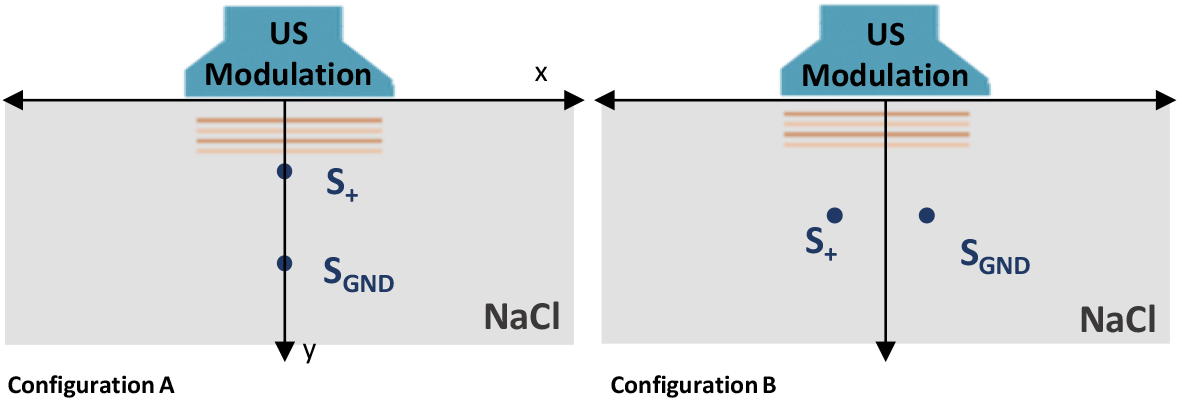
A simulated 2D conductive medium consists of NaCl (0.001mol/m3) and two polarizable metal electrodes (diameter: 0.6mm). Two lead fields are configured: US propagation direction parallel (configuration A: left panel) and perpendicular (configuration B: right panel) to the lead vector (S+ to SGND).

The ion concentration as the conservation of mass in the electrolyte phase was obtained by solving the PNP equation:

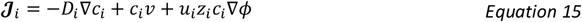

where 𝒥_*i*_ is the ion flux (SI unit: mol·m^-2^·s^-1^),*ϕ* is the electrical potential,*c*_*i*_ is the concentration, *u*_*i*_ is ion mobility, *D*_*i*_ is diffusion coefficient, *z*_*i*_ is the valence value, *i* denotes the ion species, *v* is the flow velocity.

The general Nernst-Plank equation considers three mechanisms for ion transportation:

1. Diffusion (the first term);
2. Convection/advection (the second term);
3. Electromigration (the third term).

In the proposed FE model, the convection term was neglected due to the assumption that the sodium chloride (NaCl) phantom was stationary. Thus, Equation 15 was simplified as

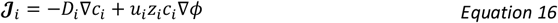

where *D*_*i*_ and *u*_*i*_ used in the model are derived from Equation 13 and Equation 14, considering the acoustic modulation of the ion hydration radius *r* and solvent viscosity *η*.

There is no established theory yet for the pressure-dependency of the ion hydration radius *r* and solvent viscosity *η* . An empirical formula was proposed for ion hydration radius *r* in our study referring to the molecular dynamics simulation data [38, 39]:

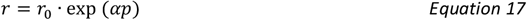

where α is the constant for pressure-dependent ion hydration radius, *p* is the acoustic pressure exported from the acoustic field simulation in the k-Wave® toolbox. A linear relationship was used to formulate the pressure-dependency of solvent viscosity *η*:

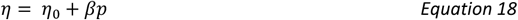

where *η*_0_ is yielded from the material function of water viscosity at room temperature in COMSOL, the constant *β* is the slope linear fitted from the experimental measurements of water viscosity at various static pressure in [40].

Consider there are no chemical reactions of the Na^+^ and Cl^-^ in the solution. The conservation of mass for both ion species satisfies

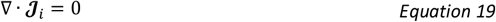

The electrochemical coupling between the electrical potential and the ion transportation in an electrolyte is described by the

Poisson equation (Gauss’s law):

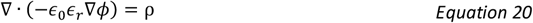

where ϵ_0_and ϵ_*r*_ are the vacuum permittivity and relative vacuum permittivity, respectively; *ρ* is the charge density (SI unit: C/m^3^), which depends on the ion concentration via:

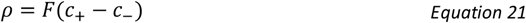

where *c*+ and*c*− are the concentration of cation and anion, respectively; *F* is the Faraday constant. The EEI interface was built based on the Gouy–Chapman–Stern model [41, 42], which predicts the diffusive layer thickness as the size of the debye length according to Equation 11.

Table 1 shows the parameters used in the FE models.

**Table 1.** Parameters for the FE Simulation.

| Characteristics | Value |
| --- | --- |
| NaCl Concentration (bulk) $c$ | 0.001 [mol/m<sup>3</sup>] |
| Electrode diameter | 0.6 [mm] |
| Standard electrode potential | 1 [V] |
| Applied electrode potential on S+ | 500 [mV] |
| US center frequency | 2.5 [MHz] |
| US peak pressure | 2.5 [MPa] |
| US pulse cycles | 4 |
| US Mechanical Index | 1.58 |
| US transducer width | 14.8 [mm] |
| Sound speed in electrolyte | 1480 [m/s] |
| Sound speed in electrode | 5500 [m/s] |
| Reference radius $r_0$ of solvated Na<sup>+</sup> | 3.58 [Å] |
| Reference radius $r_0$ of solvated Cl<sup>-</sup> | 3.32 [Å] |
| Constant for pressure-dependent ion hydration radius $\alpha$ | -0.1 [MPa<sup>-1</sup>] |
| Constant for pressure-dependent solvent viscosity $\beta$ | 0.24 [s<sup>-1</sup>] |

The radius of the solvated Na<sup>+</sup> and Cl<sup>-</sup> refer to [1]

## Results

### The Generated AE Field

Figure 6 show the simulated diffusion coefficient *D* (top row) and ion mobility *u* (bottom row) of Na^+^ and Cl^-^ when the US waves arrived at the middle region in configuration A. It reveals the local modulation of the ion diffusion coefficient *D* and mobility *u* by the acoustic waves. The magnified images present different modulation effects on the ion migration properties (*D* and *u*) from the acoustic compressional and rarefaction phases.

**Figure 6.**
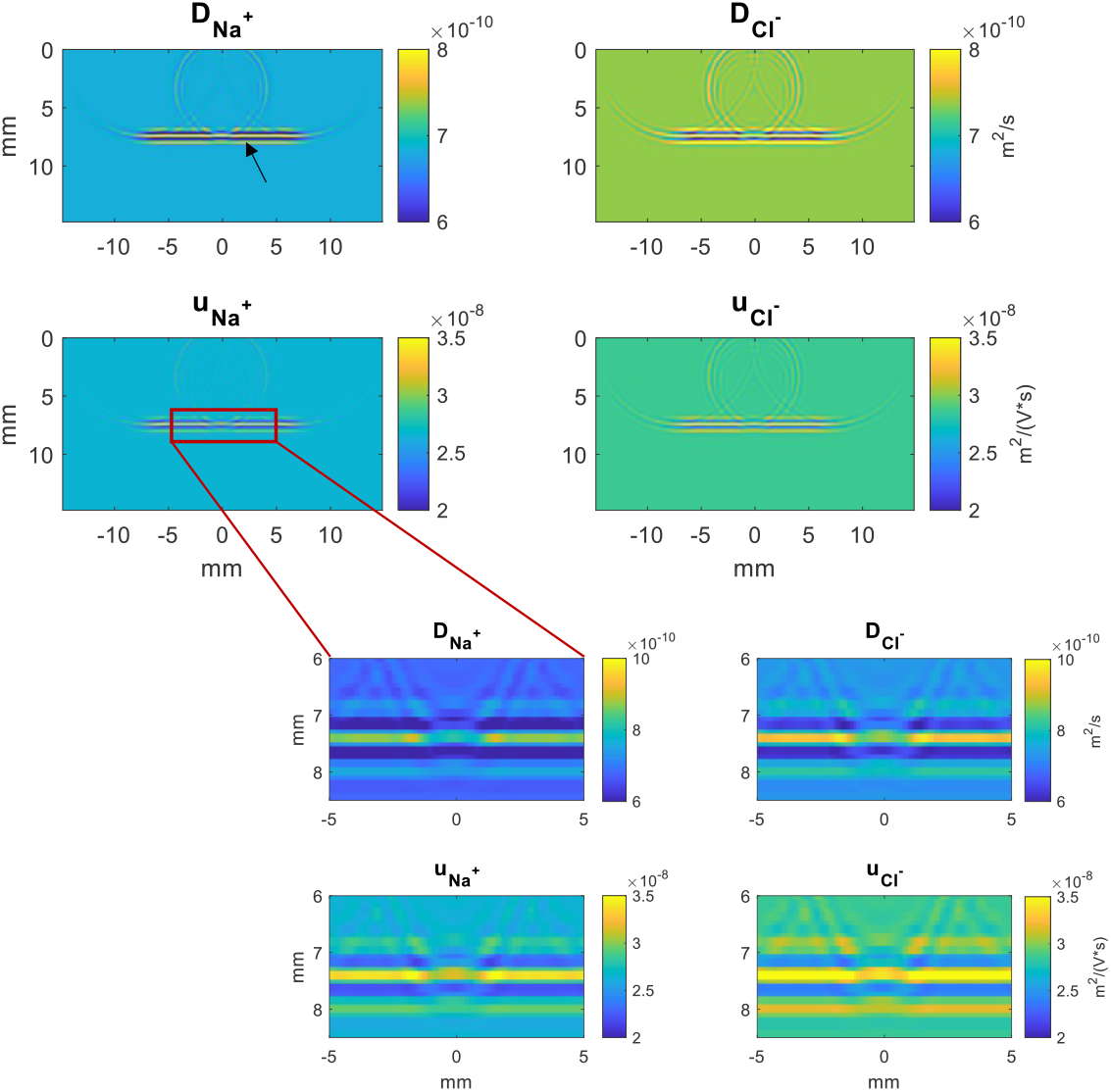
The simulated diffusion coefficient (D) and ion mobility (u) distributions of Na^+^ and Cl^-^ when the US wave arrived in the middle region in configuration A, where the lead vector is parallel to the US propagation direction. The black arrow indicates the wavefront, and the region enclosed by a red box is magnified and shown in the magnified images were taken from the spatial extent enclosed by the red rectangle.

The simulated raw AE fields were bandpass-filtered (2–3 MHz) to isolate the frequency components corresponding to the ultrasound excitation. Figure 7 show the raw (left column in each panel) and band-passed (BP: right column in each panel) electrical potential field (Φ) and the current density fields (*J*_*x*_ and *J*_*y*_) at the moment the ultrasound wave reaches the middle region in configurations A (panel A) and B (panel B), respectively. The applied background electric field originates from the +500 mV bias applied to the positive supply electrode S+ (red region in Φ map). The raw Φ and ***J*** fields collectively show the modulated fields and the background electrical field, while the BP fields extract only the oscillatory components at the insonification frequency, representing the typical AE signals measured experimentally.

**Figure 7.**
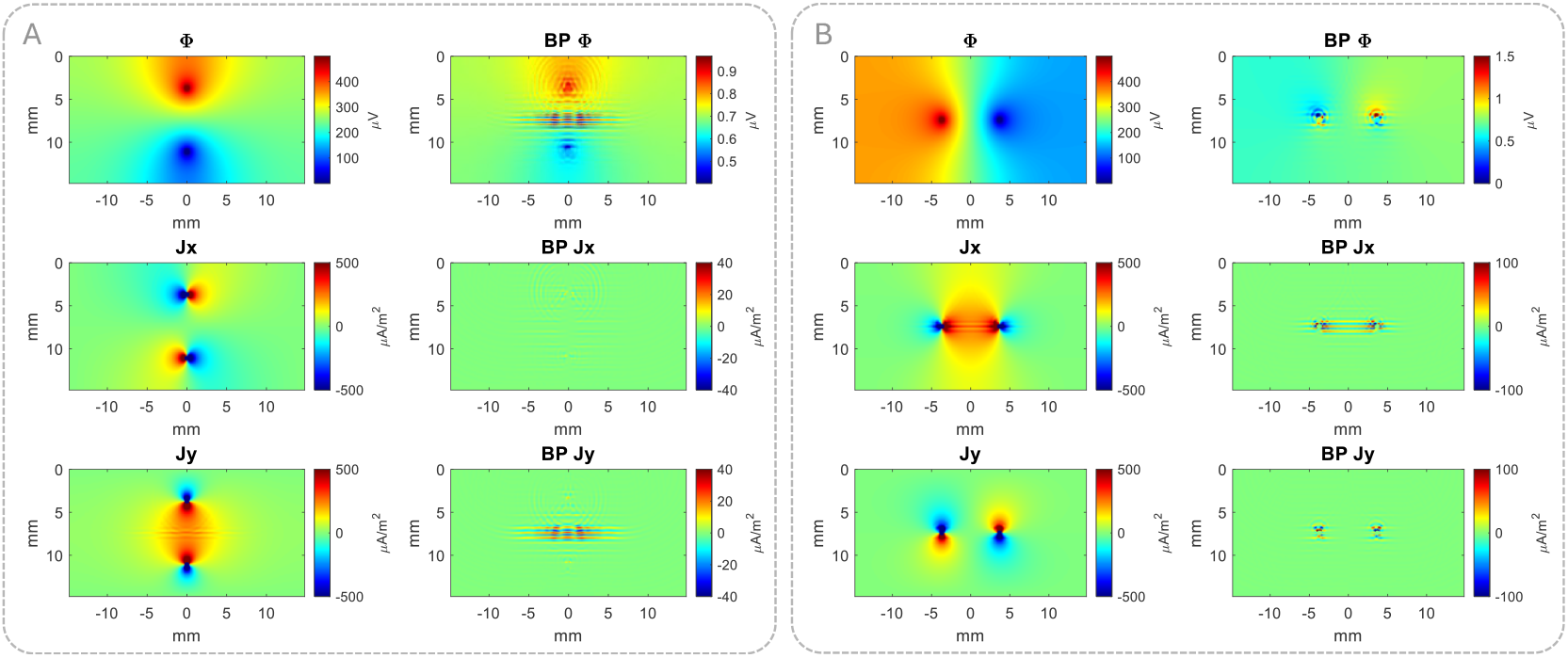
The raw and band-passed (BP) electrical potential field *ϕ* and the current density field (J_x_ and J_y_) when the acoustic waves arrived in the middle region for configuration A (left panel)and configuration B (right panel).

The *BP* Φ exhibited signal amplitudes of around 1 *μV* in both configurations, given the applied potential is 500 mV. This simulated signal magnitude aligns with the µV-scale AE signals previously recorded in NaCl experiments under 12 mA and 28 mA current excitation [3, 43].

The spatial distribution of the current density signals differed notably between the two configurations. In configuraton A, the amplitude of *BP J*_*y*_ exceeded that of *BP J*_*x*_, whereas in Configuration B *BP J*_*x*_ is larger than *BP J*_*y*_. The different signal characteristics stems directly from the change in the relative angle between the US propagation direction and the lead field vector. Besides, the signal amplitudes of the BP AE fields in configuration B were generally larger than in configuration A. This is due to the direct acoustic modulation on the electrode surface at the observation time for configuration B. The phenomenon will be further discussed in the subsequent section.

Across both configurations, the electrical potential *ϕ* and current density (*J*_*x*_ and *J*_*y*_) showed higher magnitudes around the electrode than in the bulk medium. This agreed with the spatial distribution of the simulated AE fields with dipole electrodes in 0.9% NaCl [22, 23, 44], which are based on the theory proposed in [4] via conductivity modulation. It is important to note that those earlier simulations were not designed to probe the fundamental generation mechanism of the AE effect; instead, they used the simulated AE signals to validate imaging reconstruction methods or to optimize the experimental setups. In those works, AE signals *V*_*AE*_ was computed as a volume integral of the product of the AE interaction coefficient *K*, electrical resistivity *ρ*_0_, acoustic pressure *P*, and the dot product of the recording lead field ***J***^*L*^and the source current density ***J***^*I*^:

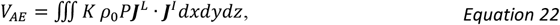

In comparison, the AE fields in our study are generated based on acoustic modulation to the ion behaviours, then coupling the resulting chemical transport with the electric field via multiphysics modelling. Given this mechanistic focus, the subsequent analysis will emphasize current density over electrical potential for two primary reasons. First, the core relationship between conductivity and ion mobility is fundamentally expressed through current density, which serves as the direct link between macro-scale measurements and micro-scale ion motion. Second, because our simulation used a voltage source, acoustic modulation of conductivity manifested primarily as a change in current. In contrast, the studies cited earlier employed current sources, where modulation appeared as a change in potential.

### Tmporal profiles of Current density

Figure 8 show the 1D temporal profiles of the current density (*J*_*x*_ and *J*_*y*_) at the central monitoring point (red cross: x=0.1*λ*, y=12.4*λ, λ*: acoustic wavelength). The blue and orange lines are the temporal profiles with the applied voltage source at 500mV and -500mV, respectively. These profiles depict how local current density at the central monitoring point was modulated as the acoustic wave propagated through (beginning around t = 12.5 *λ*/c). The modulated current density followed the insonification waveform and reversed polarity when the applied electric potential switched from 500mV to -500mV in both configurations A (upper panel) and B (lower panel). Significant direct current (DC) components were observed in the direction of the lead vector for both configurations (*J*_*y*_ for configuration A and *J*_*x*_ for configuration B). Correspondingly, the largest peak-to-peak amplitudes of the modulated signal also align with this direction. This indicates that the amplitude of the acoustically modulated signal is proportional to the current density of the existing electrical field.

**Figure 8.**
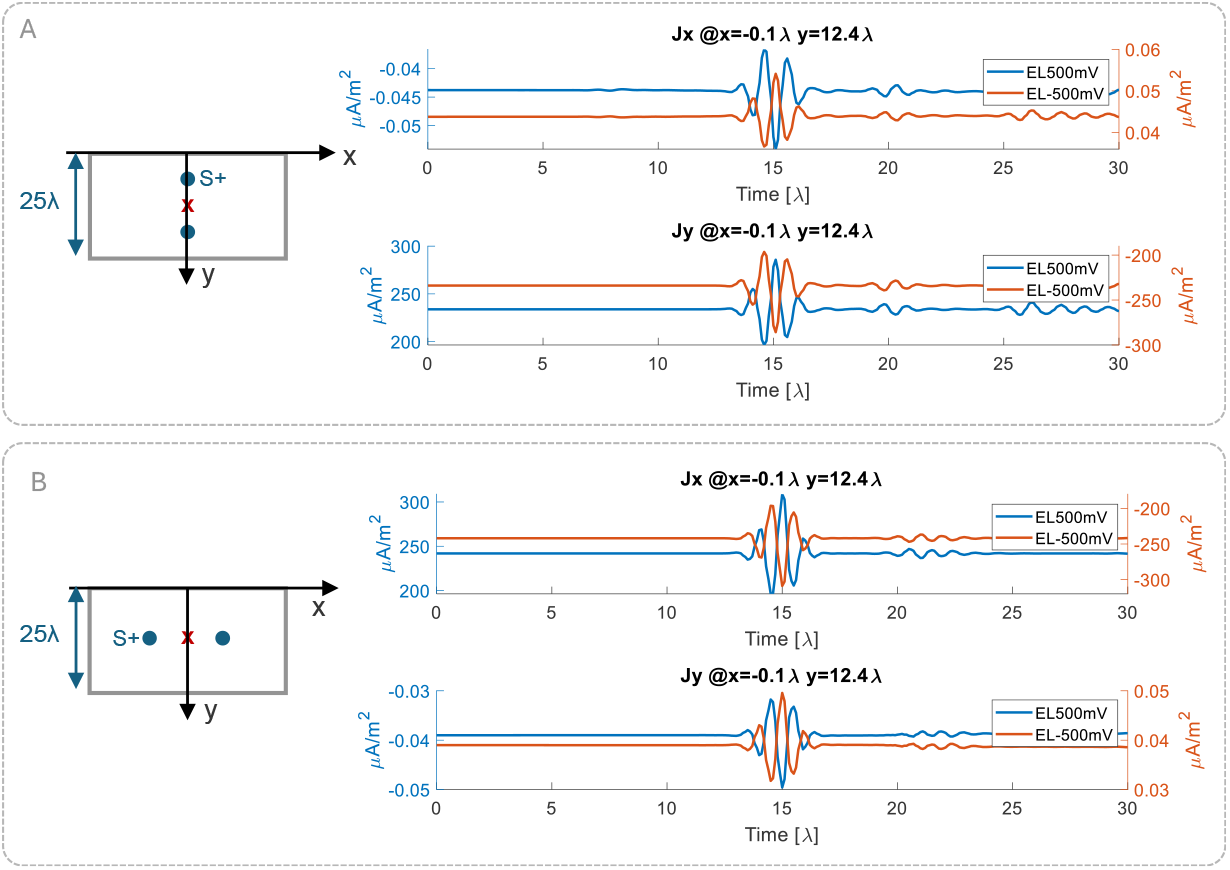
The induced local current density changes at the center monitoring point in configuration A (upper panel) and configuration B (lower panel) when the applied electrical potential on S+ is 500mV (blue line) and -500mV (orange line).

### Spatial profiles

Figure 9 shows 1D spatial profiles of the cutlines (grey arrows) in electrical potential (*ϕ*) field and current density fields (*J*_*x*_ and *J*_*y*_) for configuration A (left column) and configuration B (right column) at a fixed time point. The schematics on the top row show the location of the ultrasound wavefront at the time instance of interest, namely the arrival of the ultrasound wave at the middle region. For both configurations, the spatial profiles highlighte a pronounced modulation of the local current density inside the acoustic pulse (black arrows) than the outer area. The acoustic modulation is predominantly aligned with the lead field vector: *J*_*y*_ for configuration A (orange line in bottom left panel) and *J*_*x*_ for configuration B(yellow line in upper right panel).

**Figure 9.**
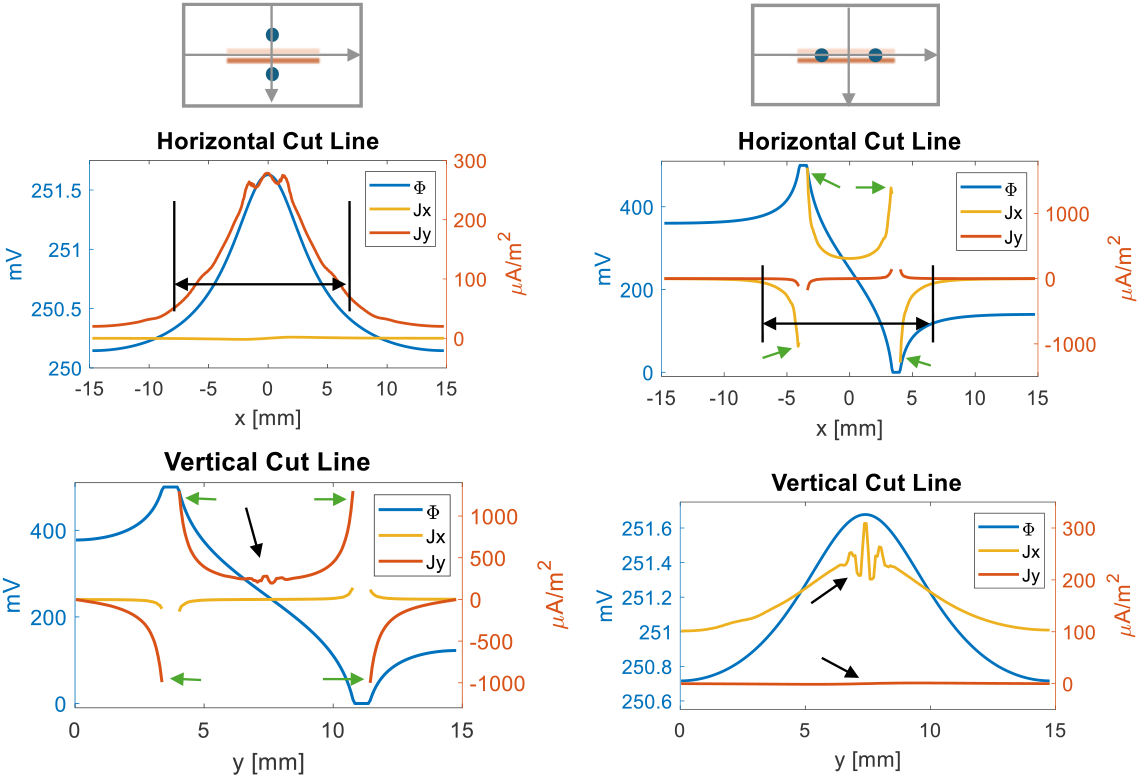
1D spatial profiles of the electrical potential and current density distribution at the time instance of the acoustic wave arrival in the middle region for configuration A (left column) and configuration B (right column). The two schematics on the top indicate the location of the ultrasound wavefront at the time instance of observation (i.e., the acoustic wave arrival in the middle region) and the spatial cutlines. Profiles are taken along these cutlines; black arrows highlight regions of acoustically modulated current.

Vertical cutline for configuration A (bottom left panel) and the horizontal cutline for configuration B (upper right panel) show the profiles through the two supply electrodes (S_+_ and S_GND_). In configuration A (bottom left panel), the electrodes were not directly modulated at the time point of observation. Thus, it revealed an inherent current flow on the electrode surface (green arrow) due to the drastic electrical potential difference in the EEI. In configuration B (upper right panel) at the time instance of observation, the incoming ultrasound waves directly act on the two supply electrodes simultaneously. The acoustic modulation of the EEI (green arrow) was observed from the cutlines of both *J*_*x*_ and *J*_*y*_.

To directly compare acoustic modulation within the bulk conductive medium to that at EEI, Figure 10 show the induced “AE signals” between two time points of observation:

**Figure 10.**
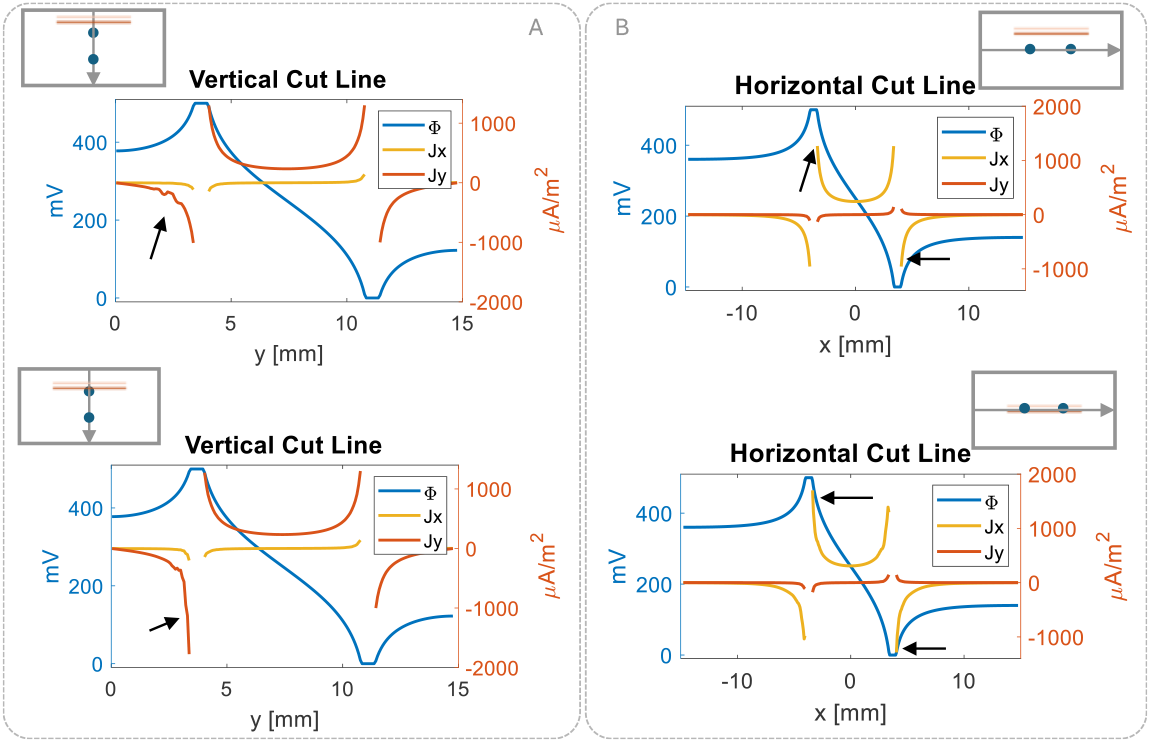
The comparison between the induced “AE signals” from the acoustic modulation of the conductive medium (upper row) and that from the acoustic modulation of the EEI (bottom row) in configuration A (left column) and configuration B (right column). The induced current density changes on EEI exhibited a larger signal amplitude (black arrow in upper row) than that from the acoustic modulation on the conductive medium near the electrodes (black arrow in bottom row).

1. when the acoustic wave was approaching the source electrode (upper panels) and
2. when the acoustic wave was in contact with the surface (bottom panels).

In configuration A (left panel) and B (right panel), a clear shift in current density (black arrows) when the acoustic wave reached the surface of S+ (bottom row), compared to the signal near the electrode before contact (upper row). The “AE signals” from the acoustic modulation of EEI (bottom rows for both panels) are notely larger than those from the acoustic modulation of the conductive medium near the electrodes (upper rows for both panels). Given that the amplitude of the modulated signals is proportional to the pre-existed current density, the large modulation signals on the EEI could be possibly attributed to the intrinsically higher current density within EEI compared to the electrolyte. Alternatively, this could also result from the acoustic modulation on the capacitance of the EDL on the EEI, where the charge and discharge of the EDL enhanced the current density changes.

## Discussion

Simulation works on the AE effect are scarce and are mostly developed either for imaging reconstruction algorithms [22, 23, 45] or applications on soft tissues on the macroscopic scale [46]. Our study introduces the first simulation model to generate AE in the context of ion behaviors in diluted strong electrolytes, thereby expanding the scope of AE effect simulations to a microscopic level.

In the classic AE phenomenology [4, 20], the modulation of electrical conductivity is attributed to bulk compression (47%), viscosity changes due to pressure (18%), and ion mobility changes against temperature (35%). This current study expands upon these findings by examining the pressure-dependency of ion mobility through ion-solvent interaction and explores the possible interplay among the three terms outlined in the classic AE phenomenology.

To investigate the acoustic modulation on electrical conductivity, the conductivity modulation ratio in this study is calculated between the conductivity under acoustic modulation *κ*_*with US*_ and the conductivity κ without an acoustic stimulus:

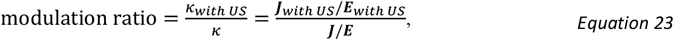

where ***J*** and ***E*** are the electrical field ***E***(*E*_*x*_, *E*_*y*_) and current density field ***J***(*J*_*x*_,*J*_*y*_), respectively.

Given the applied peak acoustic pressure *P* =2.5*MPa*, the AE coefficient *K* in this simulation was around 0.4×10^-6^ Pa^-1^, which is different than the reported impedance modulation at ∼0.9×10^-9^ Pa^-1^ for physiological solutions derived from the classic AE phenomenology in [4], and the experimental measurement of *K* in sailine at 0.03 %MPa^-1^ (0.3×10^-9^ Pa^-1^). This discrepancy arises from several factors.

First, the ion concentration in this simulation was significantly lower than that of the physiological solutions. The low ion concentration leads to the much lower conductivity used in the simulation than that of the physiological solutions. Thus, the large modulation ratio on the low conductivity in this simulation configuration still results in electrical potential changes at *μV* scale, which is comparable as reported from the physiological solutions. The results suggested the increasing conductivity modulation ratio when ion concentration is reduced. This simulation used the low ion concentration to probe ion-solvent interactions only. In other words, the solution had to be sufficiently diluted (typically <1 mol/m^3^) to ensure that ion-ion interactions could be neglected [37]. To consider the ion-ion interaction for solutions with a higher concentration, a more complex and integrated model should be developed to incorporate other theories, such as *Bjerrum theory* for ion association [37].

Second, the ion species involved in this simulation were simplified, consisting of only a single pair of cation and anion, which is fewer than those present in physiological solutions. The induced AE signals and the conductivity modulation ratio would be significantly different in the presence of ion species with various ionic strength compared to those used in this simulation.

Third, in the context of ion behavior, the current model considered the effect of ultrasonic modulation on the equilibrium among the Coulombic force, diffusive force, and frictional force, instead of regarding ultrasound modulation as an additional force that imparts an auxiliary velocity to the ions. This aspect is intrinsically linked to how fast the ions would respond to ultrasound modulation. The ionic atmosphere can be reconfigured rapidly, ranging from femtosecond to picosecond, through several collisions [47]. Such reconfiguration would be completed instantaneously, which can be encompassed by the rarefactional/compressional duration of commonly used ultrasound frequency (around microsecond for ∼MHz). This renders the ultrasound modulation of the ionic atmosphere independent of ultrasound frequency. Nonetheless, when ultrasound modulation is considered as an auxiliary force, both the duration required to attain the new mass equilibrium under ultrasound modulation and the actual exposure time render the process frequency dependent. Moreover, the direction of the auxiliary velocity from ultrasound modulation will further complicate the angle-dependency of the AE effect. To model the ion behaviors that consider both the ultrasound modulation of ion atmosphere and ultrasound-induced auxiliary velocity, a more sophisticated model, based on a more fundamental theory for Brownian motion, such as the Langevin equation, will be indispensable.

A study limitation is a lack of experimental validation for the simulation results primarily due to the instrumentation constraints. It is not feasible to isolate the ultrasound modulation on the ion atmosphere from other physical mechanisms and the parasite effects. With the advancement of instrumentation technology, the experimental measurement of the decomposed AE effect may be achieved with meticulous experimental design in the future. Another key limitation of this study is the incomplete theoretical framework regarding the pressure dependency of solvent viscosity and ion hydration radius. In the simulation, the pressure dependency of these parameters is quantified using empirical equations. Additionally, the coefficients

*α* and *β* used in these equations are fitted based on the simulated ion hydration radius and experimental measurements of water viscosity under different static pressures, respectively. Therefore, the advancement of the theory for the pressure-dependency of the ion hydration radius, and the more specific experimental measurements of the solvent viscosity under varying acoustic pressure, would further optimize the accuracy of the proposed simulation framework.

## Conclusion

AE-based functional ultrasound imaging offers unique contrast by probing the electrical properties of biological tissues. While research has progressed from phantom studies to preliminary animal experiments, AE imaging remains in its infancy. There is a pressing need for a more comprehensive theoretical model to elucidate the inherent physical processes, facilitate the analysis of signal components, and guide the design of experiments.

This work revisited the fundamental physical mechanism of the AE effect, modeled the AE generation in terms of ion-solvent interaction, and proposed a flexible FE simulation framework. A key finding is the pronounced role of the acoustic modulation on EEI, where signal generation is significantly amplified compared to modulation within the bulk electrolyte. This highlights the EEI as a critical site where local current density and double-layer dynamics enhance the acoustoelectric response. Adodpting the materials science tenet that ‘*Structure determines properties*.’, this work explores how acoustics modulate electrical properties through the modulation of the ion-solvent structures. Based on the pressure dependence of the ion atmosphere and solvent viscosity, our work highlights the value of decomposing the AE effect into its constituent ion and solvent properties. This marks a step forward in understanding these intricate interactions from a microscopic perspective and connects acoustically modulated ion dynamics to ongoing physical chemistry research. Future work is warranted to build upon these findings and develop a more complete theoretical framework.

## Notes

### Competing Interest Statement

The authors have declared no competing interest.

### Funding

- Hong Kong RGC General Research Fund: 17205022

## References

[1] E. Nightingale Jr, “Phenomenological theory of ion solvation. Effective radii of hydrated ions,” The Journal of Physical Chemistry, vol. 63, no. 9, pp. 1381–1387, 1959.

[2] P. Debye, “A method for the determination of the mass of electrolytic ions,” The Journal of chemical physics, vol. 1, no. 1, pp. 13–16, 1933.

[3] B. Lavandier, J. Jossinet, and D. Cathignol, “Experimental measurement of the acousto-electric interaction signal in saline solution,” Ultrasonics, vol. 38, no. 9, pp. 929–936, 2000.

[4] J. Jossinet, B. Lavandier, and D. Cathignol, “Impedance modulation by pulsed ultrasound,” Annals of the New York Academy of Sciences, vol. 873, no. 1, pp. 396–407, 1999.

[5] H. Zhang, and L. V. Wang, “Acousto-electric tomography.” pp. 145–149.

[6] R. Olafsson, R. S. Witte, S.-W. Huang, and M. O’Donnell, “Ultrasound current source density imaging,” IEEE Transactions on biomedical engineering, vol. 55, no. 7, pp. 1840–1848, 2008.

[7] A. Alvarez, C. Preston, T. Trujillo, C. Wilhite, A. Burton, S. Vohnout, and R. S. Witte, “In vivo acoustoelectric imaging for high-resolution visualization of cardiac electric spatiotemporal dynamics,” Applied optics, vol. 59, no. 36, pp. 11292–11300, 2020.

[8] Y. Qin, Q. Li, P. Ingram, C. Barber, Z. Liu, and R. S. Witte, “Ultrasound current source density imaging of the cardiac activation wave using a clinical cardiac catheter,” IEEE Transactions on Biomedical Engineering, vol. 62, no. 1, pp. 241–247, 2014.

[9] Q. Li, R. Olafsson, P. Ingram, Z. Wang, and R. Witte, “Measuring the acoustoelectric interaction constant using ultrasound current source density imaging,” Physics in Medicine & Biology, vol. 57, no. 19, pp. 5929, 2012.

[10] M. Allard, C. Preston, C. Huang, N.-K. Chen, and R. S. Witte, “Neuronavigation with Skull Segmentation and Acoustic Modeling for Guiding Transcranial Acoustoelectric Brain Imaging.” pp. 1–3.

[11] M. Allard, C. Preston, T. Trujillo, C. Huang, N.-K. Chen, and R. S. Witte, “MRI guided transcranial acoustoelectric imaging for safe and accurate electrical brain mapping.” pp. 1–4.

[12] C. Preston, W. S. Kasoff, and R. S. Witte, “Selective mapping of deep brain stimulation lead currents using acoustoelectric imaging,” Ultrasound in medicine & biology, vol. 44, no. 11, pp. 2345–2357, 2018.

[13] B. Berthon, P.-M. Dansette, M. Tanter, M. Pernot, and J. Provost, “An integrated and highly sensitive ultrafast acoustoelectric imaging system for biomedical applications,” Physics in Medicine & Biology, vol. 62, no. 14, pp. 5808, 2017.

[14] B. Berthon, A. Behaghel, P. Mateo, P.-M. Dansette, H. Favre, N. Ialy-Radio, M. Tanter, M. Pernot, and J. Provost, “Mapping biological current densities with ultrafast acoustoelectric imaging: Application to the beating rat heart,” IEEE Transactions on Medical Imaging, vol. 38, no. 8, pp. 1852–1857, 2019.

[15] X. Song, G. Han, Y. Zhou, M. Xu, M. Liu, and D. Ming, “A symmetrical sensor configuration for acoustoelectric brain imaging,” IEEE Sensors Journal, vol. 21, no. 20, pp. 22891–22898, 2021.

[16] Y. Zhou, X. Song, Z. Wang, F. He, and D. Ming, “Multisource acoustoelectric imaging with different current source features,” IEEE Transactions on Instrumentation and Measurement, vol. 70, pp. 1–9, 2020.

[17] Y. Zhou, X. Song, Z. Wang, X. Zhao, X. Chen, and D. Ming, “Coding biological current source with pulsed ultrasound for acoustoelectric brain imaging: application to vivo rat brain,” Ieee Access, vol. 8, pp. 29586–29594, 2020.

[18] X. Song, X. Chen, J. Guo, M. Xu, and D. Ming, “Living rat SSVEP mapping with acoustoelectric brain imaging,” IEEE Transactions on Biomedical Engineering, vol. 69, no. 1, pp. 75–82, 2021.

[19] P. Grasland-Mongrain, and C. Lafon, “Review on biomedical techniques for imaging electrical impedance,” IRBM, vol. 39, no. 4, pp. 243–250, 2018.

[20] J. Jossinet, B. Lavandier, and D. Cathignol, “The phenomenology of acousto-electric interaction signals in aqueous solutions of electrolytes,” Ultrasonics, vol. 36, no. 1-5, pp. 607–613, 1998.

[21] J. Kang, C. Huang, C. Perkins, A. Alvarez, L. Kunyansky, R. S. Witte, and M. O’Donnell, “Current source density imaging using regularized inversion of acoustoelectric signals,” IEEE transactions on medical imaging, vol. 42, no. 3, pp. 739–749, 2022.

[22] Z. Wang, and R. S. Witte, “Simulation-based validation for four-dimensional multi-channel ultrasound current source density imaging,” IEEE transactions on ultrasonics, ferroelectrics, and frequency control, vol. 61, no. 3, pp. 420–427, 2014.

[23] R. Yang, X. Li, J. Liu, and B. He, “3D current source density imaging based on the acoustoelectric effect: a simulation study using unipolar pulses,” Physics in Medicine & Biology, vol. 56, no. 13, pp. 3825, 2011.

[24] A. Hunter, “Some Observations on the Debye Effect in Electrolytes,” Proceedings of the Physical Society, vol. 71, no. 5, pp. 847, 1958.

[25] A. Rutgers, and W. Rigole, “Ultrasonic vibration potentials in colloidal solutions, in solutions of electrolytes and in pure liquids,” Transactions of the Faraday Society, vol. 54, pp. 139–143, 1958.

[26] L. Ulický, and T. J. Kemp, Comprehensive Dictionary of Physical Chemistry: E. Horwood, 1992.

[27] C. M. Brett, and O. Brett, “Principles, methods, and applications,” Electrochemistry, vol. 67, no. 2, pp. 444, 1993.

[28] P. Atkins, and J. De Paula, Elements of physical chemistry: Oxford University Press, USA, 2013.

[29] J. M. Bockris, N. Bonciocat, and F. Gutmann, “An introduction to electrochemical science,” (No Title), 1974.

[30] P. Debye, and E. Hückel, “De la theorie des electrolytes. I. abaissement du point de congelation et phenomenes associes,” Physikalische Zeitschrift, vol. 24, no. 9, pp. 185–206, 1923.

[31] M. Druchok, and M. Holovko, “Molecular dynamics study of ion hydration under pressure,” Journal of Molecular Liquids, vol. 159, no. 1, pp. 24–30, 2011.

[32] J. R. Elliott, V. Diky, I. Knotts, A. Thomas, and W. V. Wilding, “The properties of gases and liquids,” (No Title), 2023.

[33] N. Galamba, “On the effects of temperature, pressure, and dissolved salts on the hydrogen-bond network of water,” The Journal of Physical Chemistry B, vol. 117, no. 2, pp. 589–601, 2013.

[34] L. Tonti, and F. M. Floris, “How increasing pressure affects the ion hydration structure and shell properties at ambient temperature,” Journal of Molecular Liquids, vol. 328, pp. 115341, 2021.

[35] G. Jancso, K. Heinzinger, and T. Radnai, “The effect of pressure on the hydration shell of ions,” Chemical physics letters, vol. 110, no. 2, pp. 196–200, 1984.

[36] B. E. Poling, J. M. Prausnitz, and J. P. O’connell, The properties of gases and liquids: Mcgraw-hill New York, 2001.

[37] N. C. Craig, “Physical Chemistry, (Berry, R. Stephen; Rice, Stuart A.; Ross, John),” ACS Publications, 2001.

[38] A. Polidori, R. F. Rowlands, A. Zeidler, M. Salanne, H. E. Fischer, B. Annighöfer, S. Klotz, and P. S. Salmon, “Structure and dynamics of aqueous NaCl solutions at high temperatures and pressures,” The Journal of Chemical Physics, vol. 155, no. 19, 2021.

[39] R. V. Listyarini, B. M. Kriesche, and T. S. Hofer, “Characterization of the Coordination and Solvation Dynamics of Solvated Systems─ Implications for the Analysis of Molecular Interactions in Solutions and Pure H2O,” Journal of Chemical Theory and Computation, vol. 20, no. 8, pp. 3028–3045, 2024.

[40] J. Kestin, H. E. Khalifa, and R. J. Correics, “Tables of the Dynamic and Kinematic Viscosity of Aqueous NaCl Solutions in the Temperature Range 20-150 “C cand the Pressure Range 0. 1-35 MPa,” J. Phys. Chem. Ref. Data, vol. 10, no. 1, 1981.

[41] G. Gouy, “Sur la constitution de la charge electrique a la surface d’un electrolyte (electricity-on the constitution of the electrical charge at surface of an electrolyte),” Rend. C., vol. 149, pp. 654, 1909.

[42] D. L. Chapman, “LI. A contribution to the theory of electrocapillarity,” The London, Edinburgh, and Dublin philosophical magazine and journal of science, vol. 25, no. 148, pp. 475–481, 1913.

[43] R. Olafsson, R. S. Witte, and M. O’Donnell, “Measurement of a 2D electric dipole field using the acousto-electric effect.” pp. 226–235.

[44] Y. Qin, Z. Wang, P. Ingram, Q. Li, and R. S. Witte, “Optimizing frequency and pulse shape for ultrasound current source density imaging,” IEEE transactions on ultrasonics, ferroelectrics, and frequency control, vol. 59, no. 10, pp. 2149–2155, 2012.

[45] R. Yang, X. Li, A. Song, B. He, and R. Yan, “Three-dimensional noninvasive ultrasound Joule heat tomography based on the acousto-electric effect using unipolar pulses: a simulation study,” Physics in Medicine & Biology, vol. 57, no. 22, pp. 7689, 2012.

[46] X. Song, Y. Qin, Y. Xu, P. Ingram, R. S. Witte, and F. Dong, “Tissue acoustoelectric effect modeling from solid mechanics theory,” IEEE Transactions on Ultrasonics, Ferroelectrics, and Frequency Control, vol. 64, no. 10, pp. 1583–1590, 2017.

[47] D. Laage, T. Elsaesser, and J. T. Hynes, “Water dynamics in the hydration shells of biomolecules,” Chemical Reviews, vol. 117, no. 16, pp. 10694–10725, 2017.

